# Varying intestinal absorption curves of glucose and fat with different plasma insulin curves in different positions

**DOI:** 10.1101/2020.08.10.243063

**Authors:** Junhua Yang, Ling Wang, Linyang Song, Guoying Li, Xiaobao Jin, Sumin Tian

**Affiliations:** Department of anatomy, School of Biosciences & Biopharmaceutics, Guangdong Pharmaceutical University, Guangzhou, Guangdong, People’s Republic of China; Guangdong Key Laboratory of Pharmaceutical Bioactive Substances, Guangdong Pharmaceutical University, Guangzhou, Guangdong, People’s Republic of China; Class 2, Grade 2019, School of Clinical Medicine, Guangdong Pharmaceutical University, Guangzhou, Guangdong, People’s Republic of China; Department of Physiology, School of Life Sciences and Biopharmaceutics, Guangdong Pharmaceutical University, Guangzhou, China

**Keywords:** posture, intestine, absorption, metabolism, diabetes

## Abstract

It has been shown that posture could affect the rehabilitation of some diseases, and even affect the physiological metabolism and function of certain systems of the human body, including gastrointestinal absorption of glucose. Studies attributed the different gastrointestinal absorption rate in different positions to the varying rate of gastric emptying in different positions. However, it is still unknown whether the absorption rate of nutrients from the intestine varies in different positions. To verify this hypothesis, the present study was conducted using rats as subjects. After injection of glucose or chyme of safflower oil into the upper segment of jejunum of rats, curves of plasma glucose or triglyceride were drawed to evaluate and compare the potential influences on intestinal absorption by postures. We found varying intestinal absorption curves of glucose and fat with different plasma insulin curves in different positions. To be specific, the right lateral decubitus resulted in the most sharp curves, whereas the supine position the most obtuse curves with a delayed peak, both in case of glucose and fat absorption. These findings contribute to understand the position-related absorption kinetics of substances in the intestinal tract. According to this study, posture may be important for the prevention and nursing intervention of some diseases related to metabolic kinetics of glucose and lipids. It may also be important for the absorption and transport of lipophilic drugs through the mesentery lymphatic vessels.

## 1. Introduction

People spend about one third of their lives sleeping, but they seldom pay attention to the choice of sleep position. Common sleeping positions include supine, prone and lateral positions. Of them, supine position is the most used by people[1]. But probably, it is not necessarily the healthiest. When our bodies are in different positions, the organs and vessels will be in different relative positions, and the flow of body fluids will also be affected due to effect of gravity. Therefore, posture may affect the rehabilitation of some diseases, and even affect the physiological metabolism and function of certain systems of the human body.

During sleep in the supine position, the soft tissue in the mouth will fall due to gravity, which can cause double sleep apnea syndrome[2]. Moreover, when sleeping in the supine position, a large part of the heart will press the lungs, which impairs lung ventilation[3, 4]. On the contrary, sleeping in the prone position can reduce the degree of tracheal collapse and improve breathing[5]. In the prone position, the heart is more close to the chest wall, the sizes of the alveoli are more uniform, and ventilation is more smooth[4]. In addition, prone ventilation can reduce pulmonary vascular resistance and right ventricular afterload[6].Prone ventilation can also improve the ventilation function of patients with pneumonia[7]. One study has even stated that, in a sense, the longer time you spend in the prone position every day, the less damage to the lungs[8].

Moreover, side-lying, either on the left side or the right side, has been also reported to make difference with respect to several physiological or pathological aspects, including gastrointestinal absorption of glucose[9] or toxic ingestions[10], gastroesophageal reflux[1], brain glymphatic transport[11], lung function[4] and cardiac stroke volume[12].

Studies attributed the different gastrointestinal absorption rate in different positions to the varying rate of gastric emptying in different positions[9, 10]. However, it is still unknown whether the absorption rate of nutrients from the intestine varies in different positions. All the substances absorbed from the intestine flow inevitably through the veins or lymphatic vessels located in the mesentery. Anatomically, the folds and compression of the mesentery are different in different positions. Therefore, Different positions may make significant difference with respect to the absorption kinetics of intestinal nutrients.

To verify this hypothesis, the present study was conducted using rats as subjects. After injection of glucose or chyme of safflower oil into the upper segment of jejunum of rats, curves of plasma glucose or triglyceride were drawed to evaluate and compare the potential influences on intestinal absorption by postures. Glucose and lipids are drained through mesenteric veins and mesenteric lymphatic vessels, respectively. So their absorption can reflect the drainage of the two kinds of mesenteric vessels respectively.

## 2. Materials and methods

### 2.1 Animals

A total five cohort of ten-week-old male Sprague-Dawley rats were purchased from Guangdong Medical Laboratory Animal Center (Guangzhou, China). One cohort containing 20 rats were used as donors of chyme of safflower oil and the rest four cohorts (40 rats in each) were used for tests. For the first two weeks after purchased, they were housed in a specific pathogen-free conditions under 12h light-12h dark conditions with food and water available ad libitum to be allowed to accustom themselves to the new environment. Then, animals in each of the four tested cohorts were randomly divided into four groups, namely supine group (SG), prone group (PG), left-lateral-decubitus group (LG) and right-lateral-decubitus group (RG) for each specific experiment. This study was approved by the Institutional Animal Ethics Committee of Guangdong Pharmaceutical University and performed in strict accordance with the U.K. Animals (Scientific Procedures) Act, 1986.

### 2.2 Intra-jejunum injection of glucose solution

A total of 40 rats were divided randomly into four groups, 10 in each group. Rats were induced with 3% isoflurane in 100% O_2_. Once they lost the righting reflex, they were given an injection (i.p.) of ketamine (80-100 mg/kg)/xylazine (5-10mg/kg). The rats were allowed to breathe spontaneously and a heating pad was used to maintain their body temperature strictly at 37±1°C. Throughout the experiment, their heart rate, respiratory rate and oxygen saturation were continuously monitored. The anesthetic method used here was as described previously[11].

The animals were fasted overnight and then the beginning of the jejunum was ligated. A volume of 50% glucose solution (2.4 mg/g body weight) was injected into the jejunum 0.5 cm downstream of the ligation. The determination of the dose is based on previous reports[13]. The needle was injected directly into the upper jejunum with intact vasculature, as described previously[14].

After this injection, rats from SG, PG, LG and RG were immediately put individually in cages and fixed in supine position, prone position, left lateral position and right lateral position, respectively. The rats from supine group were put in cages with padding surrounding the body of the animal to preserve its position.

### 2.3 Preparation and intra-jejunum injection of chyme of safflower oil

These procedures was conducted according to a previous report[14]. In brief, a total of 20 donor 10-week-old rats were fasted about 24 hours with ample water supply before experimentation. All procedures were performed under nembutal anesthesia (30 mg/kg, Nembutal, Abbott, North Chicago, ILL). Fatty chyme was collected from the donor rats by inserting a cannula into the upper jejunum. A volume of 1 ml safflower oil was then administered by gastric intubation. The resulting chyme was rich in small clear globules of fat and was collected into ice-packed centrifuge tubes. Once the chyme from each animal was collected, it was immediately frozen at −80°C. Before given to the recipient rats, all collected chyme were armed to room temperature and immediately mixed together well.

Another 40 rats were divided randomly into four groups, 10 in each group. Rats were anaesthetized as described above (see the first paragraph of section 2.2). The animals were fasted overnight and then the beginning of the jejunum of the recipient rats was ligated. A volume of 0.5 ml prepared mixed chyme was injected into the jejunum 0.5 cm downstream of the ligation without over-distention of the wall. Then the abdominal wound was closed. Then, the rats in each of the groups were fixed in the corresponding position as described above (see the last paragraph of section 2.2).

### 2.4 Intragastric load of glucose solution

Another 40 rats were divided randomly into four groups, 10 in each group. The animals were fasted overnight and then they were anaesthetized as described above (see the first paragraph of section 2.2). A dose of 50% glucose solution (2.4mg/g body weight) was then administered by gastric intubation, as used previously[13].Then, the rats in each of the groups were fixed in the corresponding position as described above (see the last paragraph of section 2.2).

### 2.5 Intragastric load of safflower oil

Another 40 rats were divided randomly into four groups, 10 in each group. The animals were fasted overnight and then they were anaesthetized as described above (see the first paragraph of section 2.2). A volume of 1 ml safflower oil was then administered by gastric intubation. Then, the rats in each of the groups were fixed in the corresponding position as described above (see the last paragraph of section 2.2).

### 2.6 Detection of plasma glucose Level

The moment when a rat was placed in its specific position was defined as 0 min and blood was repeatedly collected from tail vein at 0, 6, 12, 30, 60 and 120 min. The time points for collecting blood were chosen according to a previous study[13]. The plasma glucose level in each sample was determined by a blood glucose meter.

### 2.7 Detection of plasma insulin

The procedure for blood collection was the same as described in section 2.6. Insulin level in each plasma sample was measured using a Rat Insulin ELISA Kit (Elabscience, Wuhan, China)

### 2.8 Detection of plasma TG

Blood was repeatedly collected from tail vein at 0, 15, 60, 120, 180 and 240 min. The time points for collecting blood were chosen according to a previous study involving intra-jejunum injection of chyme of safflower oil into small intestine and detection of TG level in tail vein[15]. TG level in each plasma sample were determined with Triglyceride Test Kit (Baiao Laibo, Beijing, China).

### 2.9 Statistical analyses

The data were statistically analyzed using the SPSS 25.0 statistical software (Chicago, IL, USA). The data are expressed as the mean ± standard deviation of the mean (SD). One-way ANOVA followed by Bonfferoni’s *post hoc* test was performed with *p* < 0.05 was considered to indicate a statistically significant difference.

## 3. Results

To specifically investigate the potential effect of position on intestinal absorption, we first carried out the intra-jejunum injection experiment using glucose and fat. After the finding of these expected effects, we then performed the intragastric load experiment. The following were what were found here. The plasma insulin curves induced by glucose absorption in both experiments were also observed.

### 3.1 Varying intestinal absorption curves of glucose with different plasma insulin curves in different positions in intra-jejunum injection experiment

As shown in Fig.1A, SG rats had a less sharp curve of plasma glucose level compared with all other groups in intra-jejunum injection experiment. LG and RG rats had the most sharp and similar curves of plasma glucose level among four groups. The trend of curve of plasma glucose level of PG rats lied between those of SG and LG/RG rats. All groups had their peaks of these curves at the same tested time point, 6 min after intra-jejunum injection of glucose. The distribution of plasma insulin curves of four groups (Fig.1C) are like those of glucose except that the SG rats’ insulin curve had a delayed peak than all other groups. Fig.2 shows the specific differences of the glucose (Fig.2A) and insulin (Fig.2B) levels among groups at each test time points.

**Fig.1.**
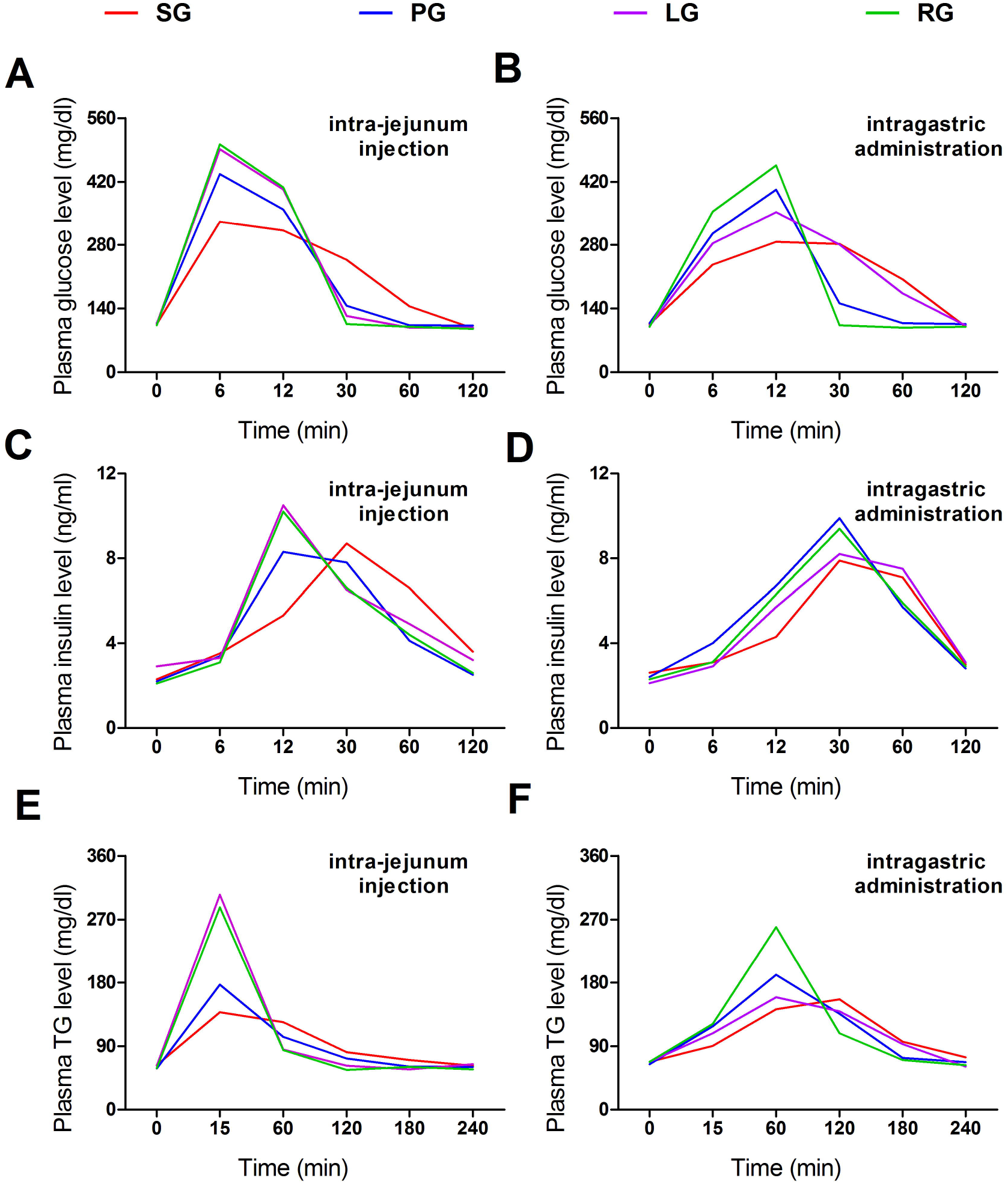
Curves of plasma levels of glucose, TG and insulin in both experiments. Each turning point in each curve represents the mean value of molecules levels in plasma collected at the corresponding detection time point in each group. *n =* 10. SG, supine group; PG, prone group; LG, left lateral decubitus group; RG, right lateral decubitus group.

**Fig.2.**
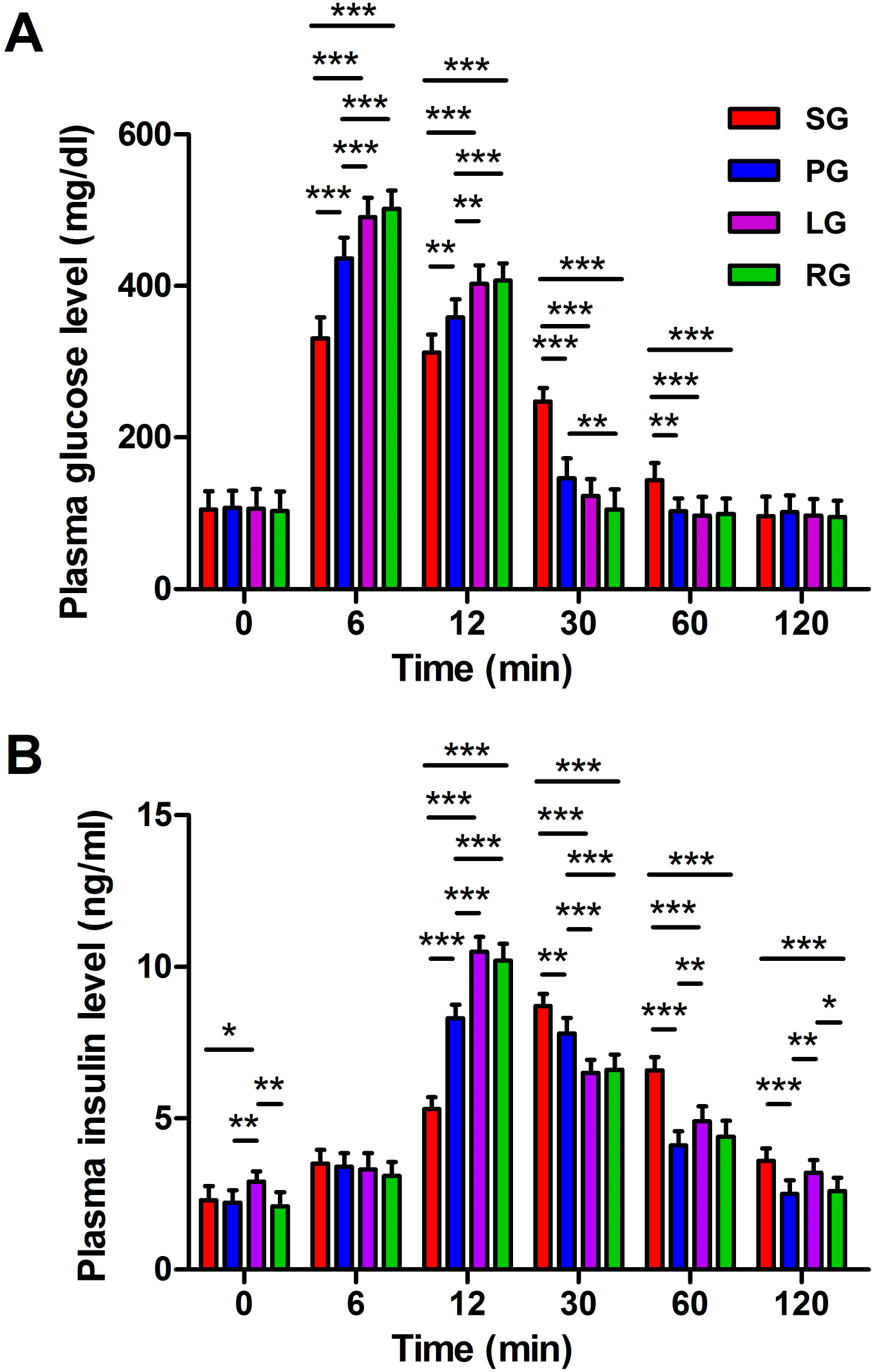
Data from intra-jejunum glucose injection experiment. **(A-B)** Bars represent mean value of glucose (A) and insulin (B) levels in plasma collected at the corresponding detection time point in each group. Data are expressed as means ± SD. One-way ANOVA followed by Bonfferoni’s post hoc test; *n =* 10; \**p* < 0.05, \*\**p* < 0.01, \*\*\**p* < 0.001. SG, supine group; PG, prone group; LG, left lateral decubitus group; RG, right lateral decubitus group.

### 3.2 Varying intestinal absorption curves of fat in different positions in intra-jejunum injection experiment

Plasma TG was detected as the index reflecting fat absorption. As shown in Fig.1E, SG rats had a less sharp curve of plasma TG level compared with all other groups in intra-jejunum injection experiment. LG and RG rats had the most sharp and similar curves of plasma TG level among four groups. The trend of curve of plasma TG level of PG rats lied between those of SG and LG/RG rats. All groups had their peaks of these curves at the same tested time point, 15 min after intra-jejunum injection of glucose. Fig.3 shows the specific differences of the TG levels among groups at each test time points.

**Fig.3.**
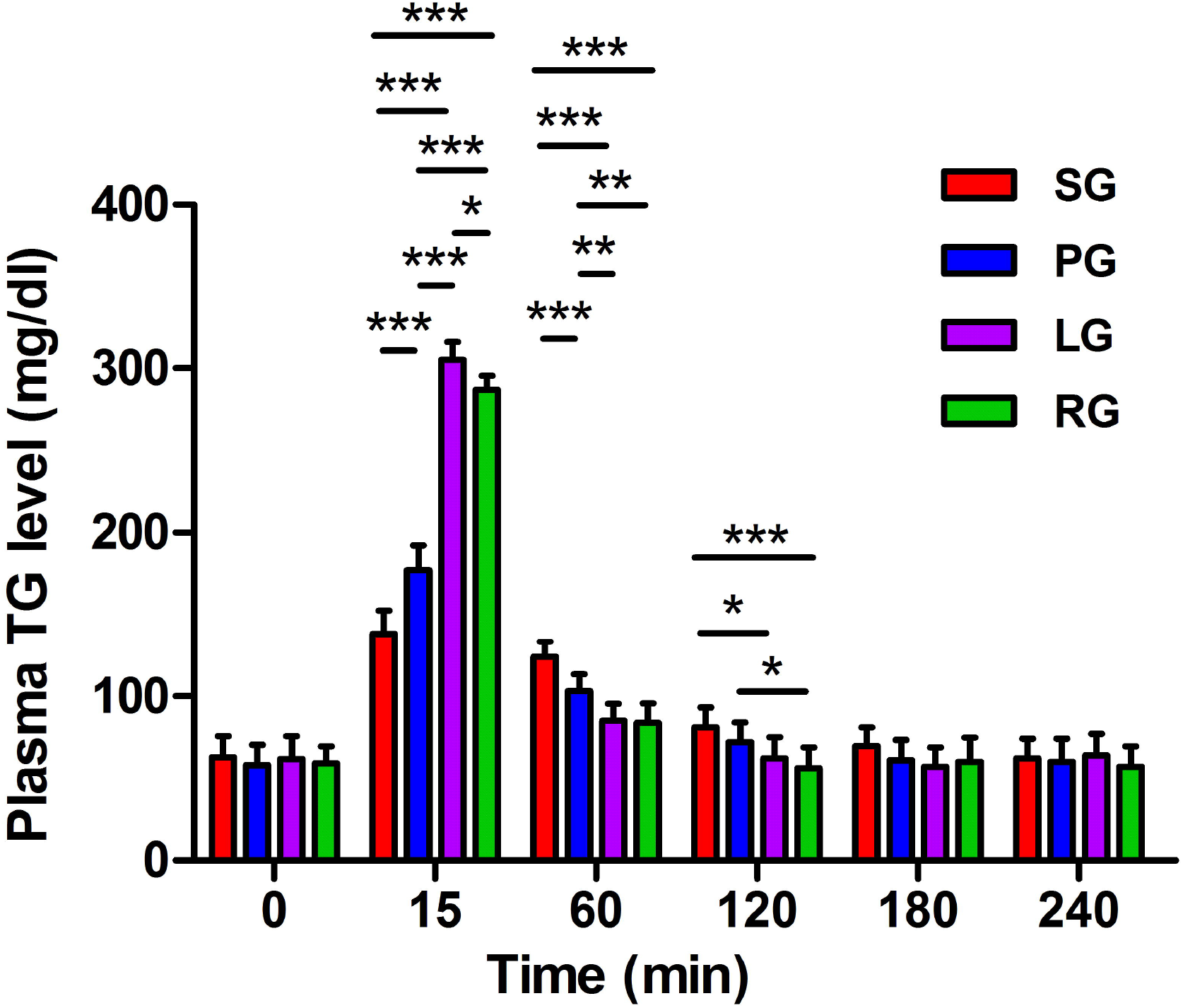
Data from intra-jejunum oil chyme injection experiment. Bars represent mean value of TG levels in plasma collected at the corresponding detection time point in each group. Data are expressed as means ± SD. One-way ANOVA followed by Bonfferoni’s post hoc test; *n =* 10; \**p* < 0.05, \*\**p* < 0.01, \*\*\**p* < 0.001. SG, supine group; PG, prone group; LG, left lateral decubitus group; RG, right lateral decubitus group.

### 3.3 Varying absorption curves of glucose in different positions in intragastric load experiment

As shown in Fig.1B, SG rats had a less sharp curve of plasma glucose level and had a delayed curve peak than all other groups in intragastric load experiment. RG rats had the most sharp curve of plasma glucose level among four groups. Compared with glucose curves in intra-jejunum injection experiment, the curves of all groups had a delayed peak with the longest delay in SG rats (to the tested time point, 30 min). Moreover, all peak values of plasma glucose levels in intragastric load experiment (Fig.1B) decreased about 50 mg/dl than those in intra-jejunum injection experiment (Fig.1A). Unlike in the intra-jejunum injection experiment, the curve of LG rats was more close to that of SG, rather than RG. The distribution of plasma insulin curves of four groups (Fig.1D) are like those of glucose (Fig.1B). Fig.4 shows the specific differences of the glucose (Fig.4A) and insulin (Fig.4B) levels among groups at each test time points.

**Fig.4.**
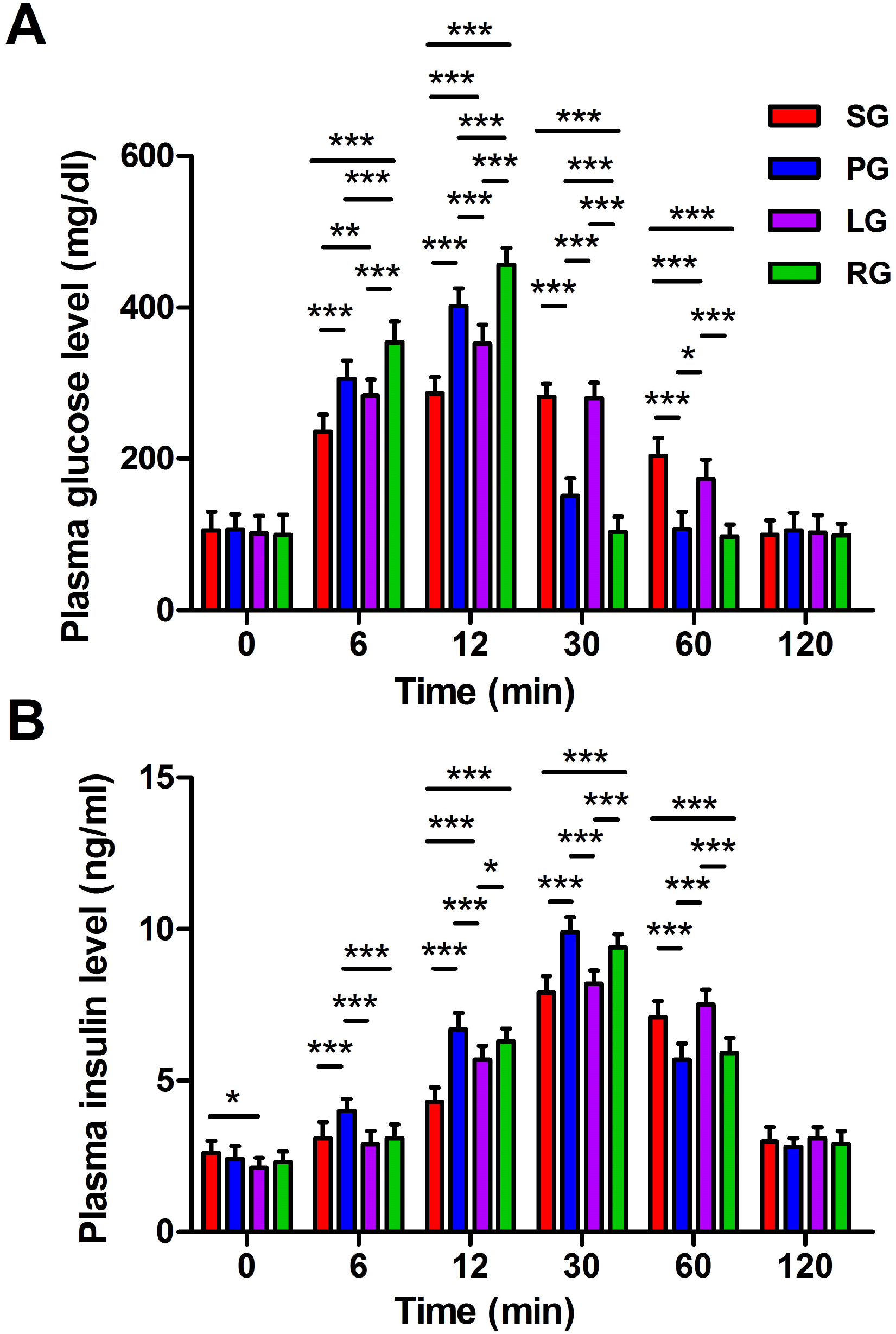
Data from in intragastric glucose load experiment. **(A-B)** Bars represent mean value of glucose (A) and insulin (B) levels in plasma collected at the corresponding detection time point in each group. Data are expressed as means ± SD. One-way ANOVA followed by Bonfferoni’s post hoc test; *n =* 10; \**p* < 0.05, \*\**p* < 0.01, \*\*\**p* < 0.001. SG, supine group; PG, prone group; LG, left lateral decubitus group; RG, right lateral decubitus group.

### 3.4 Varying absorption curves of fat in different positions in intragastric load experiment

As shown in Fig.1F, SG and LG rats had a less sharp curve of plasma TG level compared with the other groups in this intragastric load experiment. Unlike in intra-jejunum fat chyme injection experiment, only rats in RG had the most sharp curve of plasma TG level in this intragastric load experiment experiment. The trend of curve of plasma TG level of LG rats lied more close to that of SG rats. Moreover, all peak values of plasma TG levels in intragastric load experiment (Fig.1B) decreased than those in intra-jejunum injection experiment (Fig.1A). Fig.5 shows the specific differences of the TG levels among groups at each test time points.

**Fig.5.**
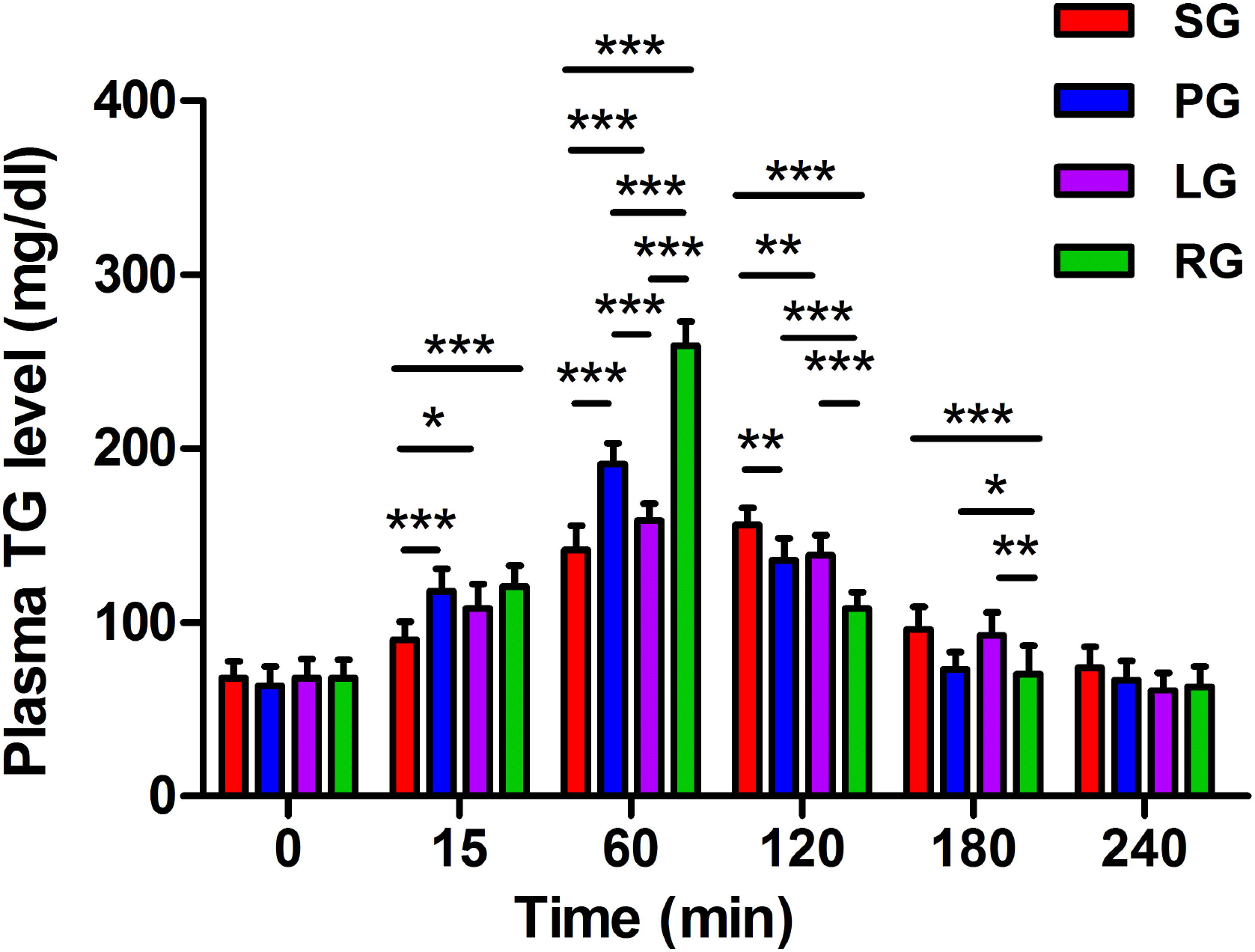
Data from intragastric oil load experiment. Bars represent mean value of TG levels in plasma collected at the corresponding detection time point in each group. Data are expressed as means ± SD. One-way ANOVA followed by Bonfferoni’s post hoc test; *n =* 10; \**p* < 0.05, \*\**p* < 0.01, \*\*\**p* < 0.001. SG, supine group; PG, prone group; LG, left lateral decubitus group; RG, right lateral decubitus group.

## 4. Discussion

This study revealed for the first time that intestinal absorption curves of glucose and fat varies according to different positions in rats. In addition, the plasma insulin curves after glucose load in upper intestine were also found different according to different positions in rats.

Either in intra-jejunum injection experiments or in intragastric load experiments, the results of this study showed that supine position negatively influenced absorption to the largest extent, while right lateral decubitus position positively influenced absorption to the largest extent. These findings suggest that the right lateral decubitus position may be the optimal position and that the supine the opposite one with respect to intestinal absorption.

The absorption curves of LG rats were obviously different in intra-jejunum injection experiment and intragastric load experiment. In the former experiments, the absorption curves of LG rats were nearly same as curves of RG rats, but they become more similar to curves of SG rats in the later experiments. This may result from the smaller rate of gastric emptying in left lateral decubitus position, as previously demonstrated[9, 10].

The pressure of lymphatic circulation is lower than that of blood circulation, so we have thought that supine position would have a greater effect on lipid absorption through the lymphatic pathway than on glucose absorption through the superior mesenteric vein. However, The experimental results do not support this idea. The possible explanation is that, unlike the portal-mesenteric vein system, the lymphatic circulation can use the valve to turn the compression of intestinal peristalsis into a power to promote lymph drainage, so that in the supine position, lymphatic drainage does not slow down significantly due to greater compression of the intestines.

In the intra-jejunum injection experiments, the absorption curves of the LG rats is very similar to that of the RG rats. This should be due to the similar degree of mesenteric extension in both lateral decubitus positions, which is larger than that in the situation of supine position. A larger extension means slighter pressure on the drainage vessels in mesenteric and less tortuosity of these vessels. The curves of PG rats lies between SG rats and rats of both lateral decubitus positions, which may be related to the press on the abdominal wall from outside and the lack of larger extension of the mesentery in the prone position.

The metabolic process of glucose and TG will undoubtedly affect their concentration in the blood. Admittedly, the rate of intestinal absorption of sugars and lipids will be more accurate if glucose levels in superior mesenteric vein blood and TG levels in thoracic duct lymph are measured. However, the required related surgical procedures will not only change the position, tension and integrity of the mesentery, but also affect the fixation of prone and both lateral recumbent positions. These change will affect the credibility of the differences in the results among the groups in this study. Therefore, this study only detected the levels of the three molecules in tail vein blood. Considering that the metabolic process of glucose and TG will affect their concentration in the blood, and this effect will vary in the presence of position-induced different absorption rates. Therefore, this study did not draw the area under the curve of each curve. Nevertheless, the absorption curve and insulin level curve presented in this study generally reflect real-time blood glucose, TG and insulin levels and their physiological fluctuations in different posture. This will help us understand the position-related absorption kinetics of substances in the intestinal tract.

The digestion and absorption of lipids is the central link of the lymphatic transport of lipophilic drugs[16]. The drugs absorbed by intestinal epithelial cells will first enter the lymphatic system without entering the liver through the portal vein, thus avoiding the metabolism of drugs into the liver. According to the experimental results of this study, the right lateral decubitus position position may promote the absorption of lipids and the transport in the lymphatic vessels, which will exert an effect on their metabolic kinetics.

As well known, many high-incidence diseases, including diabetes, hyperlipidemia and obesity, are related to abnormal insulin release and metabolic kinetics of glucose and lipids. Therefore, the findings of this study suggest that posture may be important for the prevention and nursing intervention of these diseases. People usually lie to take a nap just after lunch. Therefore, the sleeping position may have a significant effect on glucose and lipid metabolism. In addition, the conclusion of this study also provide valuable basic experimental clues as to whether long-term bedridden people need to pay attention to the choice of posture, although the exact conclusion need further clinical research to provide evidence.

## Acknowledgments

We thank Mr. Taoqi Tao (ORCID: 0000-0002-2770-9568/GDPU), Mrs. Yinyin Xie (ORCID: 0000-0002-5858-3873/GDPU) for their valuable discussions and help with this investigation.

## Author Contributions

J.Y. and S.T put forward idea, research methods and designed study program; J.Y., W.L., L.S. and S.T performed experiments and collected the data; J.Y. performed statistical analyses; J.Y. and L.S. wrote the manuscript; G.L. and X.J. reviewed and edited the manuscript. All authors read and approved the final manuscript.

